# Up-Regulation of αCaMKII Impairs Cued Fear Extinction and NMDAR-Dependent LTD in the Lateral Amygdala

**DOI:** 10.1101/2020.08.11.247270

**Authors:** Shuming An, Jiayue Wang, Xuliang Zhang, Yanhong Duan, Junyan Lv, Dasheng Wang, Huan Zhang, Gal Richter-Levin, Xiaohua Caoa

**Author notes:** These authors contributed equally to this work (Shuming An, Jiayue Wang, Xuliang Zhang).

## Abstract

Impaired fear extinction is one of the hallmark symptoms of post-traumatic stress disorder (PTSD). The roles of αCaMKII have been not extensively studied in fear extinction and LTD. Here, we found PTSD susceptible mice exhibited significant up-regulation of αCaMKII in the lateral amygdala (LA). Consistently, increasing αCaMKII in LA profoundly not only caused PTSD-like symptoms such as impaired fear extinction and anxiety-like behaviors, but also attenuated NMDAR-dependent LTD at thalamo-LA synapses, reduced GluA1-Ser845/Ser831 dephosphorylation and AMPAR internalization. Suppressing the elevated αCaMKII to normal level could completely reverse both PTSD-like symptoms and the impairments in LTD, GluA1-Ser845/Ser831 dephosphorylation and AMPAR internalization. Intriguingly, deficits in AMPAR internalization and GluA1-Ser845/Ser831 dephosphorylation were detected not only after impaired fear extinction, but also after attenuated LTD Our results demonstrate for the first time GluA1-Ser845/Ser831 dephosphorylation and AMPAR internalization are molecular links between LTD and fear extinction, and suggest αCaMKII may be a potential molecular determinant of PTSD.

## INTRODUCTION

Although some progresses have been made in understanding the molecular and cellular mechanisms of post-traumatic stress disorder (PTSD) recently, effective treatment for PTSD is still lacking. Since impaired fear extinction is one of the core symptoms of PTSD (Michopoulos et al., 2014; Yehuda et al., 2015), and fear extinction is the basis for psychological exposure therapy (M. R. Milad & Quirk, 2012), a deeper understanding of the molecular and cellular substrates underlying fear extinction would have important implications for developing the more effective treatment for PTSD.

At the synaptic level, long-term depression (LTD) has been implicated in fear extinction (Bennett, Arnold, Hatton, & Lagopoulos, 2017). N-methyl-D-aspartate (NMDA) GluN2B receptor antagonist can abolish both LTD at thalamo-lateral amygdala (T-LA) synapses and fear extinction (Dalton, Wu, Wang, Floresco, & Phillips, 2012). Moreover, deletion of kinesin superfamily proteins (KIFs) 21B impairs both hippocampal LTD and contextual fear extinction (Morikawa, Tanaka, Cho, Yoshihara, & Hirokawa, 2018). Besides, aquaporin-4 deficiency facilitates both NMDAR-dependent hippocampal LTD and fear extinction (Wu et al., 2017). Optogenetic delivery of LTD conditioning to the auditory input to LA facilitates cued fear extinction (Nabavi et al., 2014). Taken together, these findings indicate that there may be a link between LTD and fear extinction. NMDAR-dependent a-amino-3-hydroxy-5-methyl-4-isoxazolepropionic acid receptor (AMPAR) internalization is involved in fear extinction (Bai, Zhou, Wu, & Dong, 2014; Dalton, Wang, Floresco, & Phillips, 2008; J. Kim et al., 2007; Lin, Mao, Su, & Gean, 2010). Notably, disruption of AMPAR internalization impairs fear extinction (Dalton et al., 2008; J. Kim et al., 2007). Conversely, the promotion of AMPAR internalization facilitates fear extinction (Bai et al., 2014; Lin et al., 2010). It has been well known that AMPAR internalization also participates in LTD (Brebner et al., 2005; Collingridge, Isaac, & Wang, 2004). Thus, we wonder whether AMPAR internalization is a direct link between fear extinction and LTD.

At the molecular level, CaMKII is the major kinase mediating NMDAR-dependent synaptic plasticity, AMPAR trafficking and memory (Collingridge et al., 2004). In mammals, CaMKII has four isoforms, α, β, γand δ(Colbran & Soderling, 1990; Hell, 2014), and the α isoform is predominantly expressed in the forebrain (Kennedy, McGuinness, & Greengard, 1983). On the one hand, αCaMKII plays a crucial role in long-term potentiation (LTP) and memory formation (Kerchner & Nicoll, 2008; J. Lisman, Yasuda, & Raghavachari, 2012). On the other hand, αCaMKII is also required for NMDAR-dependent hippocampal LTD. For example, both CaMKII inhibitor and αCaMKII knock out could block LTD in CA1 (Coultrap et al., 2014). Moreover, αCaMKII is activated during LTD expression (J. Y. Delgado et al., 2007; Lu, Isozaki, Roche, & Nicoll, 2010). In αCaMKII-F89G transgenic (TG) mice, αCaMKII overexpression in the forebrain impairs LTD in anterior cingulate and medial prefrontal cortices, and disrupts behavioral flexibility (J. Ma et al., 2015; Wei et al., 2006). However, whether and how αCaMKII in LA affect LTD at T-LA synapses and cued fear extinction are still unknown.

To better illuminate the mechanism of cued fear extinction, thereby understanding the mechanism of PTSD, using the behavioral profiling approach (Ardi, Albrecht, Richter-Levin, Saha, & Richter-Levin, 2016), we identified PTSD susceptible mice with cued fear extinction deficit and anxiety-like behaviors from the trauma-exposed mice. It is worth noting that increased αCaMKII was detected in LA of PTSD susceptible mice. To determine whether increased αCaMKII can cause PTSD-like symptoms, we employed an inducible and reversible chemical-genetic technique to temporally and spatially manipulate αCaMKII level in the forebrain of αCaMKII-F89G TG mice, as well as using adeno-associated viral (AAV) vectors to elevate αCaMKII specifically in LA of C57BL/6J mice. Consistently, up-regulation of αCaMKII induced PTSD-like symptoms including cued fear extinction deficit and anxiety-like behaviors, which could be reversed by suppressing elevated αCaMKII to normal level. In addition, we prove that GluA1-Ser845/Ser831 dephosphorylation and AMPAR internalization are the links between cued fear extinction and NMDAR-dependent LTD at T-LA synapses.

## RESULTS

### PTSD susceptible mice exhibit increased αCaMKII and reduced AMPAR internalization in LA

PTSD susceptible individuals were identified in UWT-exposed group (23 male mice) and 4-CS/US-exposed group (23 male mice) by employing the behavioral profiling approach described in MATERIALS AND METHODS section. PTSD susceptible mice had persistently higher level of cued freeze responses through extinction trials (Fig. 1B, PS-UWT vs Control, F_(4, 85)_ = 6.33, P < 0.001; PS-4CS/US vs Control, F_(4, 85)_ = 4.70, P < 0.01), spent significantly less time in the center area of open field (OF) chamber (Fig. 1C, PS-UWT vs Control, P < 0.01; PS-4CS/US vs Control, P < 0.001), in the light zone of light/dark box (LD) test (Fig. 1D, PS-UWT vs Control, P < 0.01; PS-4CS/US vs Control, P < 0.001), and in the open arms of water zero maze (OM) test (Fig. 1E, PS-UWT vs Control, P < 0.001; PS-4CS/US vs Control, P < 0.001) compared with control mice. Behavioral profiling revealed that only 7 mice each group showed PTSD-like symptoms in 23 mice exposed to UWT or 23 mice exposed repeatedly to US/CS.

**Figure 1.**
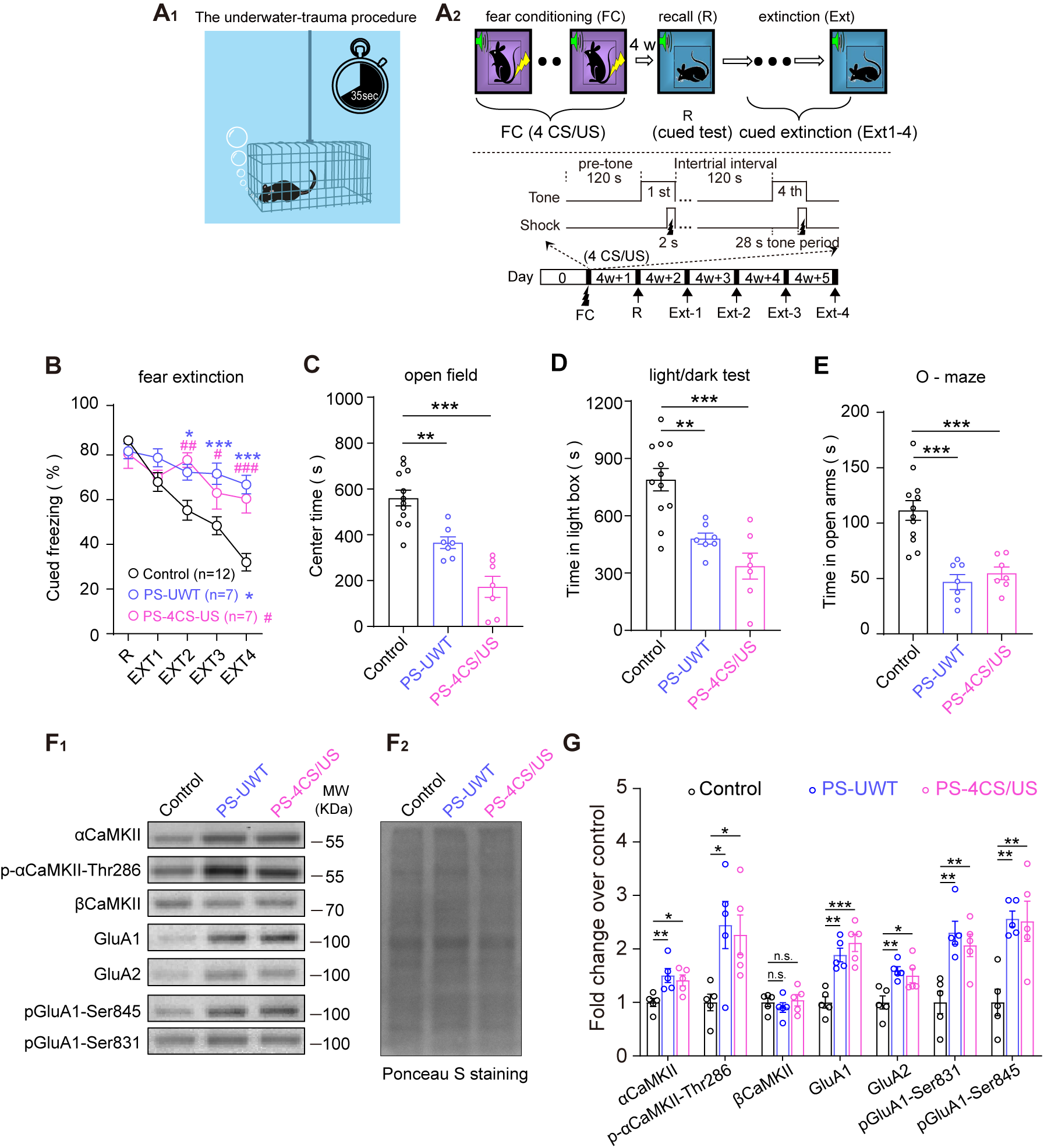
PTSD susceptible mice with cued fear extinction deficit and anxiety-like behaviors exhibited significant up-regulation of αCaMKII and down-regulation of AMPAR internalization in LA. (**A_1-2_)** Schematic illustration for identifying PTSD susceptible mice following UWT **(A_1_**, PS-UWT) or 4-CS/US pairings (**A_2_**, PS-4CS/US) exposure. (**B-E**) PTSD susceptible mice exhibited the higher level of freezing responses in the fear extinction (B), and anxiety-like behaviors in OF(C), DL(D), OM(E) tests. PTSD susceptible mice spent significantly more time freezing during extinction (B, two-way ANOVA followed by multiple comparisons with Bonferroni’s correction), less time in center area of OF chamber, in the light box of DL test and in the open arms of OM tests (C-E, one-way ANOVA followed by multiple comparisons with Bonferroni’s correction) compared to control mice (control, n = 12; PS-UWT, n = 7; PS-4CS/US, n = 7). (**F_1_**) Representative blottings of LA synaptosomal region illustrating significant higher expression in αCaMKII, p-αCaMKII-Thr286, GluA1/2, GluA1-Ser831 /Ser845 phosphorylation in PTSD susceptible mice following stress exposure, but no significant change in βCaMKII expression. (**F_2_**) Ponceau S staining was used as a loading control. (**G**) Quantifications were based on the average of independent experiment (n = 5 per group). Western blotting in “Control”, “PS-UWT” or “PS-4CS/US” groups was performed after fear extinction and all the anxiety-like behavior tests. One-way ANOVA followed by multiple comparisons with Bonferroni’s correction. n.s.: not significant, * P < 0.05, ** P < 0.01, *** P < 0.001. Error bars represent s.e.m.

LA is a key brain region for fear extinction and anxiety-like behaviors (Erlich, Bush, & Ledoux, 2012; Forster, Novick, Scholl, & Watt, 2012; Grosso, Santoni, Manassero, Renna, & Sacchetti, 2018; Jacques et al., 2019; Jihye Kim et al., 2015; J. Kim et al., 2007; Krabbe, Gründemann, & Lüthi, 2018; Mahan & Ressler, 2012; Ressler, 2010; Schafe, Doyère, & LeDoux, 2005). CaMKII has been shown to be important for memory extinction (Bevilaqua et al., 2006; Burgdorf et al., 2017; Szapiro, Vianna, McGaugh, Medina, & Izquierdo, 2003). Moreover, GluA1-Ser845/Ser831 dephosphorylation and AMPAR internalization contribute to fear extinction (Bai et al., 2014; Dalton et al., 2008; Hollis, Sevelinges, Grosse, Zanoletti, & Sandi, 2016; J. Kim et al., 2007; S. Lee et al., 2013; Lin et al., 2010; Talukdar, Inoue, Yoshida, & Mori, 2018). Thus, we investigated levels of CaMKII, GluA1-Ser845/Ser831 phosphorylation and synaptic GluA1/2 expression in LA of PTSD susceptible mice and found αCaMKII and the phosphorylated (p)-αCaMKII at Thr286 (p-αCaMKII-Thr286) were significantly up-regulated in PTSD susceptible mice experienced either UWT or 4-CS/US exposure (Fig. 1FG, PS-UWT vs Control, αCaMKII, P < 0.01, p-αCaMKII-Thr286, P < 0.05; PS-4CS/US vs Control, αCaMKII, P < 0.05, p-αCaMKII-Thr286, P < 0.05). However, no significant difference was observed in βCaMKII among the three groups (Fig. 1F, PS-UWT vs Control, P > 0.05; PS-4CS/US vs Control, P > 0.05). In addition, PTSD susceptible mice had a significant higher synaptic expression levels in the synaptic GluA1/2 expression and phosphorylated GluA1-Ser845/Ser831 (Fig. 1F, PS-UWT vs Control, GluA1: P < 0.01, GluA2: P < 0.01, GluA1-Ser831: P < 0.01, GluA1-Ser845: P < 0.01; PS-4CS/US vs Control, GluA1: P < 0.001, GluA2: P < 0.05, GluA1-Ser831: P < 0.01, GluA1-Ser845: P < 0.01). Taken together, these results suggest that PTSD susceptible mice display the significantly higher level of αCaMKII, the lower level of GluA1-Ser845/Ser831 dephosphorylation and AMPAR internalization in LA.

### Increasing αCaMKII in LA is sufficient to cause PTSD-like phenotypes in both αCaMKII-F89G TG and AAV-αCaMKII mice

To further investigate whether elevated αCaMKII in LA cause PTSD-like phenotypes such as impaired fear extinction and anxiety-like behaviors, we temporally and spatially manipulated αCaMKII overexpression in αCaMKII-F89G TG mice by employing an inducible and reversible chemical-genetic technique described in MATERIALS AND METHODS section. The higher level of αCaMKII and normal morphology in LA were observed in TG mice (Supplemental information, Fig. S1).

Then, cued fear memory recall and cued fear extinction were measured after only 1-CS / US for cued fear conditioning (Fig. 2A). Given that forebrain αCaMKII overexpression impairs fear memory retrieval in our previous study (Cao et al., 2008), to examine the effect of αCaMKII overexpression on cued fear extinction in TG mice, we designed the “normal αCaMKII level during cued fear memory retrieval but elevated αCaMKII level during cued fear extinction period” paradigm by a single i.p. injection of NM-PP1 into both TG mice and WT littermates 15 mins before the first recall test of cued fear memory (Fig. 2A). Under this paradigm, TG mice exhibited normal retrieval of cued fear memory in comparison to that of wild-type littermate (Fig. 2B, TG + i.p. vs WT + i.p., P > 0.05). However, during cued fear extinction trials, as shown in Fig. 2B, a significant declining freezing behavior was observed in WT mice but not in TG mice (Fig. 2B, TG + i.p. vs WT + i.p., F _(3, 264)_ = 10.73, P < 0.001). A *post hoc* analysis revealed that TG mice exhibited significantly higher level of freezing response to the CS in cued fear extinction trial 2, 3 and 4 (Fig. 2B, TG + i.p. vs WT + i.p., P < 0.05), suggesting that elevated αCaMKII may impair cued fear extinction. In addition, TG mice spent significantly less time (Fig. 2C–E, TG + i.p. vs WT + i.p., P < 0.001) in the center area of OF chamber (Fig. 2C), in the light zone of LD test (Fig. 2D), and in the open arms of elevated plus maze (EPM) test (Fig. 2E) compared with WT mice. Together, it indicates that increased αCaMKII in LA may cause PTSD-like phenotypes.

**Figure 2.**
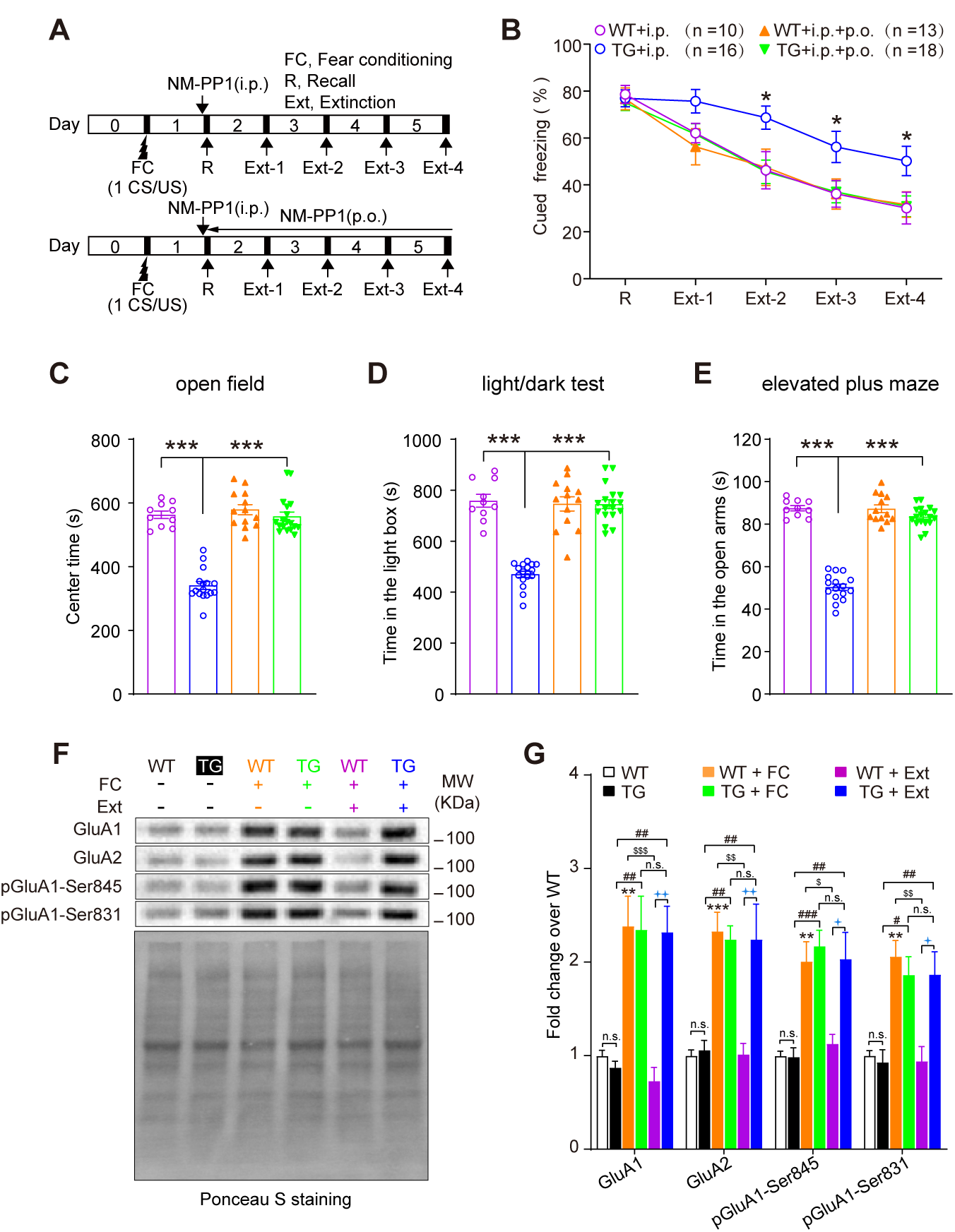
αCaMKII-F89G TG mice exhibited PTSD-like behaviors and impairments in AMPAR internalization. **(A)** The schematic of behavioral procedure for cued fear conditioning and extinction trials.**(B)** Impaired cued fear extinction in TG mice (two-way ANOVA followed by multiple comparisons with Bonferroni’s correction). Intraperitoneal (i.p.) injection and oral (p.o.) administration with NM-PP1 could rescue the cued extinction deficits of TG mice (two-way ANOVA followed by multiple comparisons with Bonferroni’s correction). (**C-E**) The higher level of anxiety-like behaviors in TG mice in the OF(C), DL(D) and EPM(E) tests after cued fear conditioning and extinction (one-way ANOVA followed by multiple comparisons with Bonferroni’s correction). (**F**) Up: Representative blottings of LA synaptosomal fractions illustrating an increase in GluA1/2, phosphorylation level of GluA1-Ser845/Ser831 in both WT and TG mice after cued fear conditioning. Down: Ponceau S staining was used as a loading control. A decrease in GluA1/2, phosphorylation level of GluA1-Ser845/Ser831 in WT mice, but not in TG mice after cued fear extinction (n = 5 per group). **(G)** Quantifications were based on the average of independent experiment. Western blotting in “WT/TG + FC” or “WT/TG + Ext” groups was performed after fear conditioning or fear extinction following with anxiety-like behavior tests, respectively (one-way ANOVA followed by Bonferroni’s multiple comparisons test). n.s.: not significant, * P < 0.05, ** P < 0.01 and *** P < 0.001 versus WT group; $ P < 0.05, $$ P < 0.01 and $$$ P < 0.001 versus WT + FC group; # P < 0.05, ## P < 0.01 and ### P < 0.001 versus TG group; 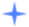 P < 0.05 and 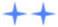 P < 0.01. Error bars represent s.e.m.

To further confirm whether PTSD-like phenotypes in TG mice are due to the overexpression of αCaMKII-F89G protein, we then designed the “normal αCaMKII level during both fear memory recall and extinction period” paradigm by i.p. injection of NM-PP1 15 min before recall test and oral (p.o.) administration throughout the entire fear extinction period (Fig. 2B). Under this “normal αCaMKII level during both fear memory recall and extinction period” paradigm, TG mice had similar freezing response with that in WT mice during cued extinction trials (Fig. 2B, TG + i.p. + o.p. vs WT + ip, P > 0.05), suggesting impaired cued fear extinction was rescued by NM-PP1 treatment in TG mice. Moreover, NM-PP1 had no effect on cued fear extinction in WT mice (Fig. 2B, WT + i.p. + o.p. vs. WT + i.p., P > 0.05), excluding the possibility that the rescuing effects by NM-PP1 were due to ‘facilitating extinction’ effects. In addition, TG mice with NM-PP1 treatments spent comparable amounts of time (Fig. 2C–E, TG + i.p. + p.o. vs. WT + i.p., P > 0.05) in the center area of OF chamber (Fig. 2C), in the light box of LD test (Fig. 2D) and in EPM test (Fig. 2E) compared with WT mice. Furthermore, TG mice without any treatment exhibited normal locomotor activity, exploratory behavior and pain threshold (Supplemental information, Fig. S2). Taken all together, we conclude that increased αCaMKII indeed is sufficient to produce PTSD-like phenotypes including impaired fear extinction and anxiety-like behaviors.

To further examine whether increasing αCaMKII specifically in LA is also sufficient to cause PTSD-like phenotypes, we bilaterally injected viral vectors AAV-αCaMKII (pAAV-TRE-αCaMKII-P2A-EGFP-CMV-rTA) into LA of C57BL/6J mice to overexpress αCaMKII specifically in LA (Fig. 3A). As expected, both αCaMKII and p-αCaMKII-Thr286 expression levels significantly increased in LA of AAV-αCaMKII mice (Fig. 3DE, αCaMKII, P < 0.001; p-αCaMKII-Thr286, P < 0.001). 24 h after 1-CS/US pairing, we performed cued fear memory test. AAV-αCaMKII mice exhibited impairment of cued fear memory during recall test (Fig. 3C, P < 0.001), which is consistent our previous finding that αCaMKII overexpression impairs the retrieval of fear memory (Cao et al., 2008). In addition, AAV-αCaMKII mice showed significantly impaired fear extinction (Fig. 3C, AAV-αCaMKII vs AAV-control, F_(4, 81)_ = 2.63, P < 0.05). A post hoc analysis revealed that AAV-αCaMKII mice exhibited the significant higher freezing responses than AAV-control mice on the 4th extinction trial (P < 0.05). In addition to deficits in cued fear extinction, AAV-αCaMKII mice showed anxiety-like behaviors (data not shown). These data suggest that elevated αCaMKII expression specifically in LA is also sufficient to result in PTSD-like symptoms.

**Figure 3.**
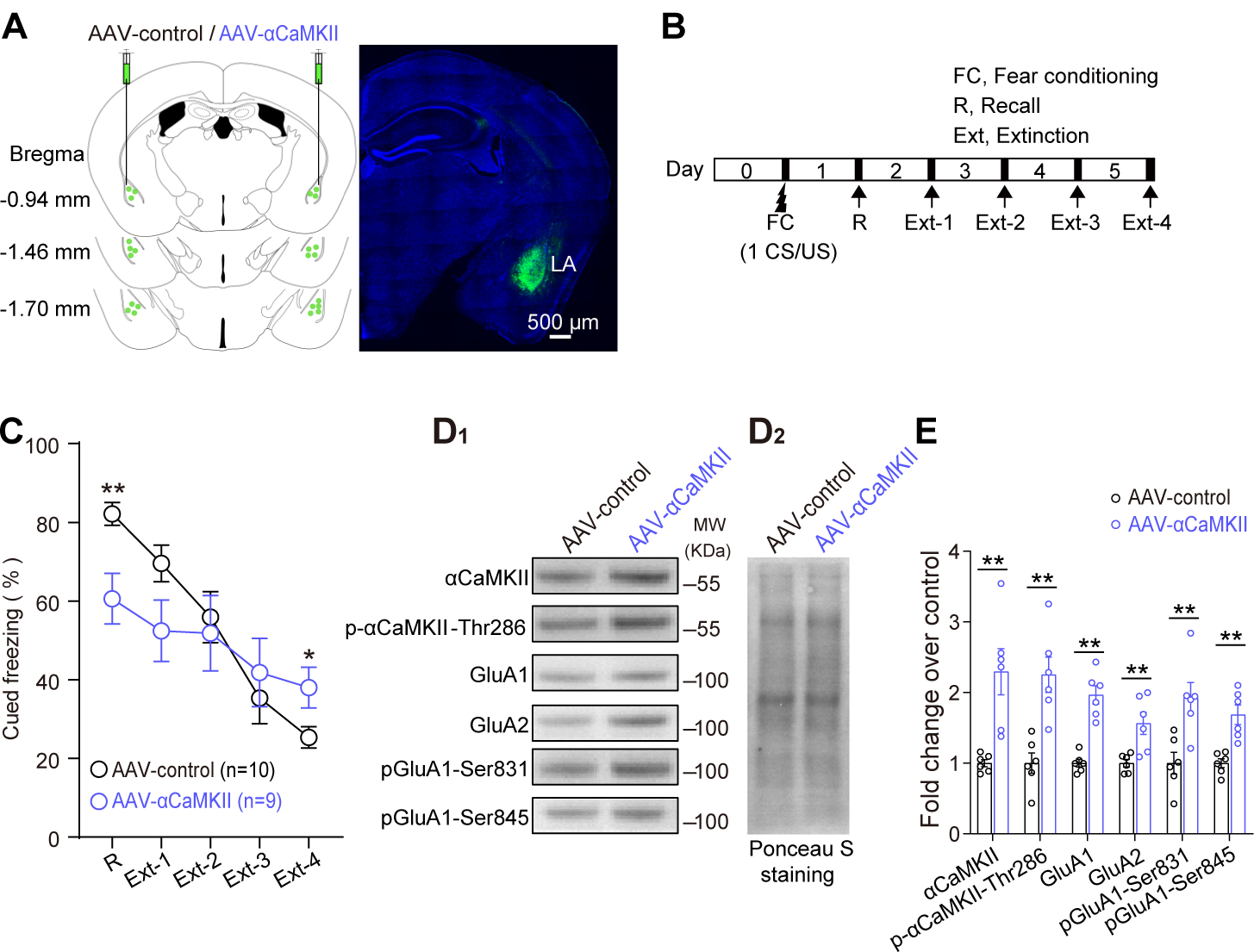
Increasing αCaMKII specifically in LA impaired the cued fear extinction and AMPAR internalization in AAV-αCaMKII mice. **(A)** Images of coronal brain slice showing the expression of eGFP (green-colored) 6 weeks after bilateral injections of pAAV-TRE-αCaMKII-P2A-EGFP-CMV-rTA virus into LA. Numbers indicate coordinates relative to bregma. Scale bar, 500 µm. **(B)** The schematic of behavioral procedure for cued fear extinction trials. **(C)** Elevating αCaMKII in LA could impair cued fear memory and fear extinction (two-way ANOVA followed by multiple comparisons with Bonferroni’s correction). (**D_1_**) Representative blottings of LA synaptosomal fractions illustrating an increased expression of αCaMKII, p-αCaMKII-Thr286, GluA1/2, phosphorylated GluA1-Ser845/Ser831 in LA of AAV-αCaMKII mice than that in AAV-control mice. (**D_2_**) Ponceau S staining was used as a loading control. (**E**) Quantifications were based on the average of independent experiment (n = 6 per group). Western blotting was performed after fear extinction and all the anxiety-like behavior tests. Statistical differences were evaluated with Student’s t test. * P < 0.05, ** P < 0.01, *** P < 0.001 Error bars represent s.e.m.

### Increasing αCaMKII in LA impairs AMPAR internalization and GluA1-Ser845/Ser831 dephosphorylation after cued fear extinction in both αCaMKII-F89G TG and AAV-αCaMKII mice

We quantified the expression of synaptic AMPAR composition subunits (GluA1/2) and GluA1-Ser845/Ser831 phosphorylation in LA before/after cued fear conditioning and extinction trials. After cued fear extinction trials, compared with cued fear conditioning trial, significant decreases in the GluA1/2 synaptic expression and GluA1-Ser845/Ser831 phosphorylation levels could be found only in WT mice (Fig. 2FG, WT + FC vs WT + Ext, GluA1: P < 0.001; GluA2: P < 0.01; pGluA1-Ser845: P < 0.05; pGluA1-Ser831: P < 0.01), but not in TG mice (TG + FC vs TG + Ext, GluA1/A2, pGluA1-Ser845/831: P > 0.05). Furthermore, the GluA1/2 synaptic expression and phosphorylated GluA1-Ser845/Ser831 in TG mice were significantly higher than that in WT mice after cued fear extinction trials (Fig. 2FG, TG + Ext vs WT + Ext, GluA1/A2: P < 0.01; pGluA1-Ser845/Ser831: P < 0.05).

Moreover, consistent with the above western blotting data from αCaMKII-F89G TG mice, synaptic GluA1/2 expression, phosphorylated GluA1-Ser845/Ser831 levels were significantly higher in LA of AAV-αCaMKII mice than that in AAV-control mice after cued fear extinction trials (Fig. 3DE, GluA1/2, pGluA1-Ser845/Ser831, P < 0.01). Taken all together, these results indicate that increasing αCaMKII specifically in LA disrupts GluA1-Ser845/Ser831 dephosphorylation and AMPAR internalization, consequently may impair cued fear extinction in both αCaMKII-F89G TG and AAV-αCaMKII mice.

### Increasing αCaMKII impairs NMDAR-dependent LTD at T-LA synapses and NM-PP1 can recover the impairments

To investigate the cellular mechanism of impaired cued fear extinction, we measured the basal synaptic transmission and synaptic plasticity at T-LA synapses in TG mice. No significant difference was observed in input-output curves, synaptic and total GluA1/2 expression of LA (Supplemental information, Fig. S3AB, TG vs WT, P > 0.05), paired-pulse depression (PPD) and synapsin expression (Fig. S3CD, TG vs WT, P > 0.05) in LA between TG and WT mice. Moreover, either tetanic or theta burst stimulations induced similar level of LTP at T-LA synapses (Fig. 4AB, TG vs WT, P > 0.05). These results indicate that αCaMKII overexpression does not affect basal synaptic transmission and LTP at T-LA synapses.

**Figure 4.**
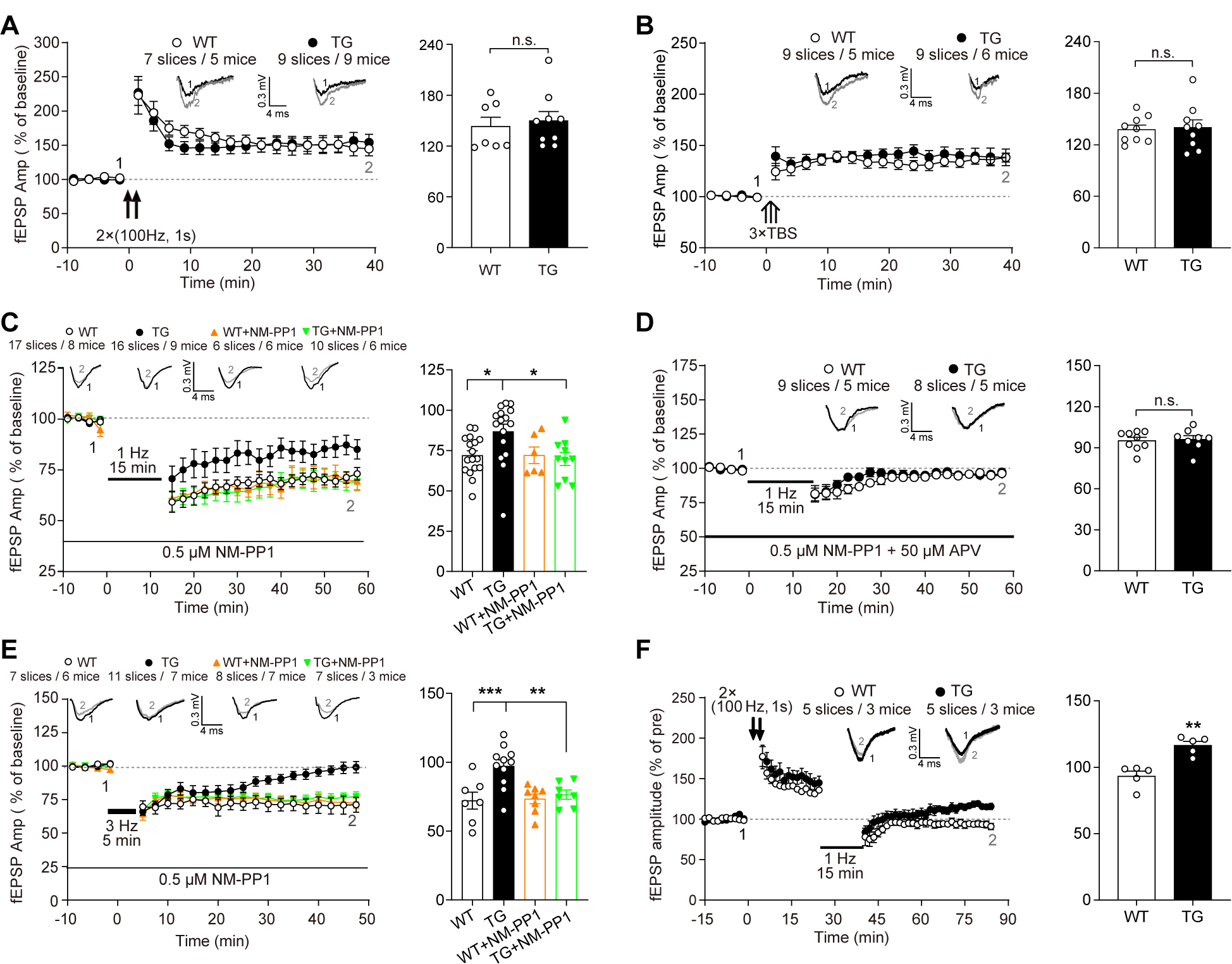
Increasing αCaMKII impairs NMDAR-dependent LTD at T-LA synapses of αCaMKII-F89G TG mice and NM-PP1 can rescue the impairments. **(A)** Similar LTP induced by high frequency stimulations (2 trains of 100 Hz stimulation for 1 s, 10 s interval) in TG slices and WT slices. In this and the subsequent figures, insets show sample traces taken at baseline (1) and the last 10 min recording (2). **(B)** Normal LTP induced by three trains of theta burst stimulations (TBS, each train consisted of 10 bursts delivered at 5 Hz, each burst consisted of 4 pulses at 100 Hz) in TG slices. **(C)** Significantly weaker LTD induced in TG slices than that in WT slice after 1 Hz (15 min) stimulation. NM-PP1 (0.5 μM) recovered the reduced LTD in TG slice to normal level. **(D)** LTD was abolished in WT and TG slices exposed to both NM-PP1 (0.5 μM) and APV (50 μM). The solid line shows the duration of both NM-PP1 and APV application. **(E)** Strong LTD could be induced by 3 Hz (5 min) stimulation in WT slices but not in TG slices, NM-PP1 (0.5 μM) rescued the impaired LTD in TG slice. **(F)** Impaired depotentiation can be observed in TG slices. All of the bar graph summarizing data obtained during last 10 min recording. Statistical differences were evaluated with Student’s t test (A, B, D and F) and one-way ANOVA followed by Bonferroni’s multiple comparisons test (C and E). n.s.: not significant, * P < 0.05, ** P < 0.01, *** P < 0.001. All values are mean ± s.e.m.

We then analyzed the effects of the αCaMKII overexpression on LTD at T-LA synapses. 1Hz-LTD in TG slices was significantly reduced (Fig. 4C, TG vs WT, P < 0.05) compared to that of WT slices, which could be recovered by 0.5 μM NM-PP1 (Fig. 4C, TG + NM-PP1 vs TG, P < 0.05), while 1Hz-LTD in WT slices was not affected (Fig. 4C, WT + NM-PP1 vs WT, P > 0.05). Notably, LTD at the T-LA synapses could be blocked by application of APV (50 μM) and NM-PP1 (0.5 μM) (Fig. 4D, TG vs WT, P > 0.05), suggesting the LTD at the T-LA synapses is NMDAR-dependent. Besides, 3Hz-LTD was blocked in TG slices (Fig. 4E; TG vs WT, P < 0.001), which could also be recovered by NM-PP1 (Fig. 4E, TG + NM-PP1 vs TG, P < 0.01), while 3Hz-LTD in WT slices was not affected (Fig. 4E, WT + NM-PP1 vs WT, P > 0.05). Likewise, TG mice exhibited deficits in the depotentiation at T-LA synapses (Fig. 4F, TG vs WT, P < 0.01). In summary, our results show that αCaMKII overexpression impairs NMDAR-dependent LTD and depotentiation at T-LA synapses in TG mice.

### Increasing αCaMKII impairs AMPAR internalization and GluA1-Ser845/Ser831 dephosphorylation during NMDAR-dependent LTD and NM-PP1 can rescue the impairments

Besides the low-frequency stimulation (LFS), brief NMDA exposure can chemically induce NMDAR-dependent LTD (H. K. Lee, K. Kameyama, R. L. Huganir, & M. F. Bear, 1998). In TG slices, NMDA application (30 μM, 3 min) could elicit a significantly weaker LTD at T-LA synapses than that in WT slices (Fig. 5A, TG + NMDA vs WT + NMDA, P < 0.01). Furthermore, 0.5 μM NM-PP1 could rescue the reduced NMDA-induced LTD in TG slices to normal level (Fig. 5A, TG + NMDA + NM-PP1 vs WT + NMDA, P > 0.05; TG + NMDA + NM-PP1 vs TG + NMDA, P < 0.01), but had no detectable effects on NMDA-induced LTD in WT slices (Fig. 5A, WT + NMDA + NM-PP1 vs WT + NMDA, P > 0.05). These results suggest that increasing αCaMKII in LA attenuates NMDAR-dependent chem-LTD at T-LA synapses in TG mice.

**Figure 5.**
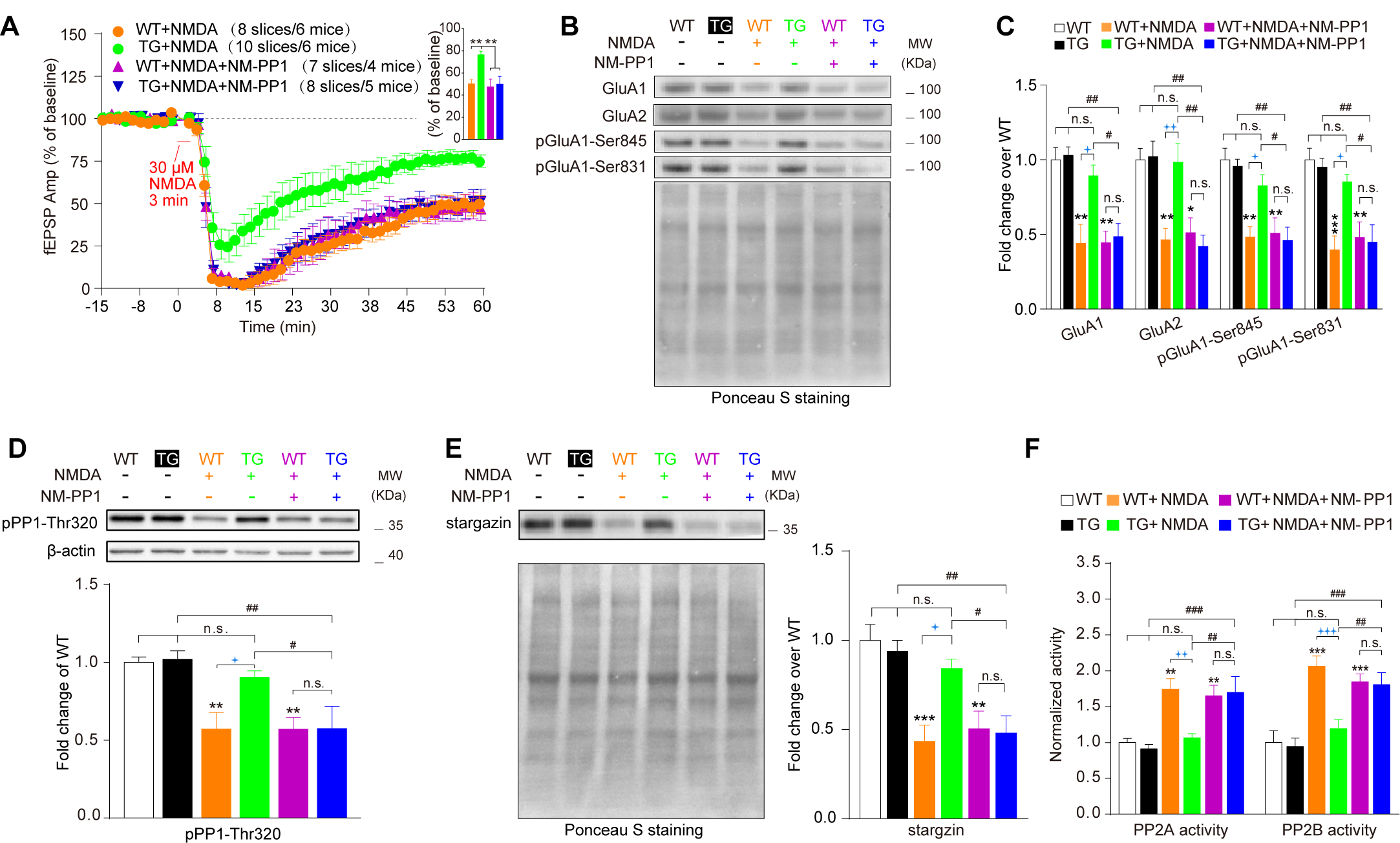
Increasing αCaMKII impairs AMPAR internalization / dephosphorylation, reduces protein phosphotase (PP) activity, and increases stargazin expression during NMDAR-dependent LTD and NM-PP1 can rescue all impairments. **(A)** Attenuated chem-LTD induced by 30 μM NMDA for 3 min in TG slices. This deficit could be rescued by 0.5 μM NM-PP1. Right-up panel: bar graph summarizing data obtained during last 10 min recording in the different groups depicted. The following Western blotting was performed 1 hour later after NMDA application. (**B**) Representative blottings of LA synaptosomal fractions illustrating a reduction in GluA1/2, phosphorylation level of GluA1-Ser845/831 in WT slices after NMDA treatment but not in TG slices. NM-PP1 could rescue these deficits in TG slices. Down: Ponceau S staining was used as a loading control. (**C)** Quantifications were based on the average of independent experiments (n = 5 per group). **(D)** Up: Representative blottings of LA synaptosomal fractions illustrating a reduction in phosphorylation level of pPP1-Thr320, indicating an increase in PP1 activity in WT mice after NMDA treatment but not in TG. NM-PP1 rescued such deficit in TG mice. Down: Quantifications were based on the average of independent experiments (n = 4 per group). **(E)** A remarkably higher level of stargazin in amygdala synaptosomal fractions in TG slices than that in WT slices after NMDA application, NM-PP1 rescued the deficit in TG mice (n = 5 per group). Down: Ponceau S staining was used as a loading control. **(F)** An increased activity of PP2A and PP2B in WT slices were exhibited after NMDA application but not in TG slices, and NM-PP1 rescued these deficits in TG mice (n = 4 per group). Statistical differences were evaluated with one-way ANOVA followed by multiple comparisons with Bonferroni’s correction. n.s.: not significant, * P < 0.05, ** P < 0.01 and *** P < 0.001 versus WT group; # P < 0.05, ## P < 0.01 and ### P < 0.001 versus TG group; 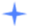 P < 0.05, 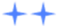 P < 0.01 and 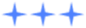 P < 0.001. Error bars represent s.e.m.

NMDA-induced LTD could elicit more widespread depression of synapse strength and share the similar molecular mechanisms to LFS-LTD such as AMPAR internalization and GluA1-Ser845/Ser831 phosphorylation (Jary Y. Delgado et al., 2007; He, Lee, Song, Kanold, & Lee, 2011; Kollen, Dutar, & Jouvenceau, 2008; H.-K. Lee, K. Kameyama, R. L. Huganir, & M. F. Bear, 1998). To investigate the molecular mechanisms underlying deficit in NMDAR-dependent LTD in TG mice, we examined the amount of some synaptic proteins after NMDA-induced LTD. NMDA application significantly decreased the GluA1/2 synaptic expression and GluA1-Ser845/Ser831 phosphorylation in WT slices (Fig. 5BC, WT+NMDA vs WT, GluA1, GluA2, pGluA1-Ser845: P < 0.01; pGluA1-Ser831: P < 0.001), but not in TG slice (Fig. 5BC, TG+NMDA vs TG, GluA1/A2, pGluA1-Ser845/831: P > 0.05). Besides, the synaptic expression of GluA1/2 and GluA1-Ser845/Ser831 phosphorylation in LA of TG slices were significantly higher than that in LA of WT slices (Fig. 5BC, TG + NMDA vs WT + NMDA, GluA1: P < 0.05; GluA2: P < 0.01; pGluA1-Ser845/Ser831: P < 0.05).Furthermore, NM-PP1 (0.5 µM) successfully rescued the impairments of AMPAR internalization and GluA1-Ser845/Ser831 dephosphorylation of LA in TG slices (Fig. 5BC, TG+NMDA+NM-PP1 vs TG, GluA1/A2, pGluA1-Ser845/831: P < 0.01), with no effect on that of WT slices (Fig. 5BC, WT+NMDA+NM-PP1 vs WT+NMDA, GluA1/A2, pGluA1-Ser845/831: P > 0.05). Collectively, it indicates that αCaMKII overexpression leads to impairment of AMPARs internalization and dephosphorylation in LA, which consequently impairs NMDAR-dependent LTD at T-LA synapses.

### Increasing αCaMKII reduces protein phosphotase (PP) activitiy and enhances stargazin expression during NMDAR-dependent LTD and NM-PP1 can recover the abnormalities

Activation of protein phosphatase 1 (PP1) contributes to LTD formation (Isabelle M. Mansuy & Shirish Shenolikar, 2006; Mauna, Miyamae, Pulli, & Thiels, 2011). Moreover, stargazin can be dephosphorylated by PP1 to induce the clathrin-dependent AMPAR endocytosis during NMDAR-dependent LTD (Bats, Groc, & Choquet, 2007; Matsuda et al., 2013). Dephosphorylation of the Thr320 residue on the C-terminal domain of PP1 can enhance PP1 activity during NMDAR-dependent LTD (Dohadwala et al., 1994; Goldberg et al., 1995). Therefore, we investigated PP1-Thr320 phosphorylation (pPP1-Thr320) and stargazin expression in LA fractions of WT and TG slices with NMDA treatment. With NMDA exposure, significant reductions of pPP1-Thr320 and stargazin expression of LA could be found only in WT (Fig. 5DE, WT+NMDA vs WT, pPP1-Thr320: P < 0.01; stargazin: P < 0.001) but not in TG slices (TG+NMDA vs TG, pPP1-Thr320, stargazing: P > 0.05). Moreover, the PP1-Thr320 phosphorylation and stargazin expression in LA of TG slices were dramatically higher than that in WT slices (Fig. 5DE, TG + NMDA vs WT + NMDA, pPP1-Thr320, stargazing, P < 0.05), suggesting that the PP1 activity and stargazin expression were abnormal in TG mice during NMDA-induced LTD. Furthermore, NM-PP1 could recover the abnormalities in PP1 activity and stargazin expression in LA of TG slices (Fig. 5DE, TG + NMDA + NM-PP1 vs TG, pPP1-Thr320: P < 0.01; stargazing: P < 0.01; TG + NMDA + NM-PP1 vs TG + NMDA, pPP1-Thr320: P < 0.05, stargazing: P < 0.05) but not affecting that of WT slices (Fig. 5DE, WT + NMDA + NM-PP1 vs WT + NMDA, pPP1-Thr320: P > 0.05; stargazing: P > 0.05).

Protein phosphatase 2A (PP2A) and calcineurin (PP2B) play important roles in LTD maintenance and induction (Pi & Lisman, 2008; Winder & Sweatt, 2001). A significant augment of PP2A/2B activity could be found in LA of WT slices (Fig. 5F, WT+NMDA vs WT, PP2A: P < 0.01; PP2B: P < 0.001), but not in LA of TG slices during LTD formation (Fig. 5F, TG+NMDA vs TG, PP2A, PP2B: P > 0.05). Besides, PP2A/2B activity was dramatically lower in LA of TG slices than that of WT slices (Fig. 5F, TG + NMDA vs WT + NMDA, PP2A: P < 0.01; PP2B: P < 0.001), during NMDA-induced LTD. Furthermore, NM-PP1 (0.5 µM) could also recover PP2A/2B activity down-regulation in LA of TG slices (Fig. 5F, TG + NMDA + NM-PP1 vs TG, PP2A, PP2B: P < 0.001; TG + NMDA + NM-PP1 vs TG + NMDA, PP2A, PP2B: P < 0.01) without affecting that of WT slices (Fig. 5F, WT + NMDA + NM-PP1 vs WT + NMDA, PP2A, PP2B: P > 0.05). Taken together, all these results suggest that αCaMKII overexpression can weaken PP1, PP2A/2B activity and increase stargazin expression in LA fractions during NMDAR-dependent LTD, which may be potential mechanisms of AMPAR internalization and NMDAR-dependent LTD impairments.

## DISCUSSION

In the present study, we reveal that PTSD susceptible mice exhibits significant up-regulation of αCaMKII, down-regulation of GluA1-Ser845/Ser831 dephosphorylation and AMPAR internalization in LA. Consistently, increasing αCaMKII specifically in LA can cause PTSD-like phenotypes such as fear extinction deficit and anxiety-like behaviors, and impairs AMPAR internalization and dephosphorylation, NMDAR-dependent LTD and depotentiation at T-LA synapses. Furthermore, deficits in AMPAR internalization and dephosphorylation are observed not only after impaired cued fear extinction *in vivo*, but also after attenuated NMDA-induced LTD in TG slices *in vitro*. Additionally, the deficits in AMPAR internalization and dephosphorylation are due to down-regulation of PP1/2A, PP2B activity and increased stargazin in TG mice. Importantly, NM-PP1, a specific inhibitor of the exogenous αCaMKII-F89G, could rescue the above deficits in αCaMKII-F89G TG mice. These data suggest up-regulation of αCaMKII may weaken activity of PP1/2A and PP2B, increase stargazing, thereby impairing AMPAR internalization and dephosphorylation, which consequently impairs LTD and fear extinction.

### αCaMKII and memory extinction

CaMKII has been shown to play an important role in the extinction of different memories. Pharmacological inhibition of CaMKII by KN-62 blocked the extinction of step-down passive avoidance performance (Bevilaqua et al., 2006; Szapiro et al., 2003). Similarly, α/βCaMKII inhibitor KN93 significantly attenuated the extinction of cocaine conditioned place preference (Burgdorf et al., 2017). Furthermore, partial reduction of αCaMKII function due to the T286A^+/–^ mutation impaired the extinction of contextual fear and spatial memories (Kimura, Silva, & Ohno, 2008). On the contrary, reduction of αCaMKII by phosphorylation at serine 331 in LA enhances cocaine memory extinction (Rich et al., 2016). Besides, increased activation of CaMKIIα in the CPEB3-knockout hippocampus reduced the extinction of spatial memories (Berger-Sweeney, Zearfoss, & Richter, 2006; Huang, Chao, Tsai, Chung, & Huang, 2014). In our study, we found that mouse models of PTSD with cued fear extinction deficit exhibited significant up-regulation of αCaMKII in LA. Furthermore, increasing αCaMKII in LA can cause PTSD-like phenotypes including impaired cued fear extinction.

### The causal relationship between elevated αCaMKII and impaired LTD

CaMKII is a major kinase mediating AMPAR trafficking and NMDAR-dependent synaptic plasticity (Collingridge et al., 2004). Specifically, CaMKII can phosphorylate AMPA receptors GluA1 subunits at Ser845/Ser831, which can promote the integration of new AMPA receptors at the postsynaptic density (Barria, Muller, Derkach, Griffith, & Soderling, 1997), further enhancing synaptic transmission. On the contrary, CaMKII has been found to interact with Arc/Arg3.1 gene product to weaken synapses by promoting AMPA internalization (Okuno et al., 2012). Recently, CaMKII has been also shown to phosphorylate GluA1 subunits at Ser567 site to promote P2X2-mediated AMPAR internalization and drive synaptic depression (Pougnet et al., 2016). In our study, we found that PTSD susceptible mice with blocked fear extinction exhibited significantly higher αCaMKII, lower GluA1-Ser845/Ser831 dephosphorylation and lower AMPA internalization in LA. To investigate whether elevated αCaMKII led to PTSD-like symptoms including impaired fear extinction, changed NMDAR-dependent LTD, GluA1 dephosphorylation and AMPA internalization in LA, we then up-regulated αCaMKII expression in αCaMKII-F89G TG and AAV-αCaMKII infected mice. We found that αCaMKII overexpression in LA caused impairments in GluA1-Ser845/Ser831 dephosphorylation, AMPA internalization, NMDAR-dependent LTD at T-LA synapses and cued fear extinction in TG mice, which could be completely rescued by a specific inhibitor (NM-PP1) of exogenous αCaMKII-F89G. These results suggest there is causality between up-regulated αCaMKII and impaired GluA1-Ser845/Ser831 dephosphorylation, defective AMPA internalization, NMDAR-dependent LTD and cued fear extinction.

### The molecular links between LTD and fear extinction

NMDAR mediates both LTD and fear extinction (Bai et al., 2014; Brebner et al., 2005; Dalton et al., 2008; Fox, Russell, Titterness, Wang, & Christie, 2007; Radulovic, Ren, & Gao, 2019). It has been reported that a GluR2-derived peptide (Tat-GluR23Y) blocked AMPAR internalization and impaired NMDAR-dependent LTD both *in vitro* (Bai et al., 2014; Brebner et al., 2005; Dalton et al., 2008) and *in vivo* (Fox et al., 2007). Moreover, NMDA NR2B receptors antagonist (Ro25-6981) blocked AMPAR internalization and disrupted fear extinction (J. Kim et al., 2007). Conversely, systemic administration of d-serine enhanced both AMPAR internalization and fear extinction (Bai et al., 2014). In addition, GluA1-Ser845/Ser831 dephosphorylation also played important roles in NMDAR-dependent LTD (Diering, Heo, Hussain, Liu, & Huganir, 2016) and fear extinction (Hollis et al., 2016; Talukdar et al., 2018). Although the above findings indicate AMPAR internalization and dephosphorylation may be links between fear extinction and LTD, supporting evidence is still lacking. In our current study, deficits in GluA1-Ser845/Ser831 dephosphorylation and AMPAR internalization were observed not only after impaired cued fear extinction *in vivo*, but also after attenuated NMDA-induced LTD in αCaMKII-F89G TG slices *in vitro*. Furthermore, a specific inhibitor of the exogenous αCaMKII-F89G (NM-PP1) could completely rescue the deficits in cued fear extinction, NMDA-induced LTD, GluA1-Ser845/Ser831 dephosphorylation and AMPAR internalization. Thus, our data demonstrate that deficits in Ser845/GluA1-Ser831 dephosphorylation and AMPAR internalization by elevated αCaMKII are molecular links between impaired NMDAR dependent-LTD and fear extinction. In other words, we demonstrate for the first time that GluA1-Ser845/Ser831 dephosphorylation and AMPAR internalization are molecular links between NMDA dependent-LTD and fear extinction.

### How does excessive αCaMKII impair AMPAR internalization and dephosphorylation during NMDAR-dependent LTD?

LTD formation requires PPs (PP1, PP2A and PP2B) activation (Kameyama, Lee, Bear, & Huganir, 1998; H. K. Lee et al., 1998). Activated PPs dephosphorylate GluA1-Ser845/Ser831 (Hu, Huang, Yang, & Xia, 2007; I. M. Mansuy & S. Shenolikar, 2006; Winder & Sweatt, 2001), which cause a reduction of open probability or conductance for AMPAR channels and finally contribute to LTD formation. Specifically, PP1 is activated through a Ca^2+^-PP2B-I1 pathway and has a more predominant role in depressing potentiated synapses, whereas PP2A is activated through PP2B/PP1 cascade or pathways independent on PP2B and mainly depresses naive synapses (Winder & Sweatt, 2001). However, in αCaMKII-F89G TG mice, αCaMKII overexpression could exhibit higher potency in the competition with PP2B for Ca^2+^/CaM, which might decrease the accessibility of PP2B to Ca^2+^/CaM and inhibit the activity of the PP2B-I1-PP1 pathway, thereby inhibiting PP1 activity. In addition, high concentration of phosphorylated αCaMKII could saturate the dephosphorylation ability of PP1, and thereby weaken PP1 dephosphorylating GluA1-Ser845 or GluA1-Ser831 (Hu et al., 2007; H. K. Lee, Barbarosie, Kameyama, Bear, & Huganir, 2000; J. E. Lisman & Zhabotinsky, 2001; I. M. Mansuy & S. Shenolikar, 2006; Winder & Sweatt, 2001). Unlike PP1, PP2A can be directly inactivated by CaMKII through phosphorylating its B’ α subunits (Fukunaga et al., 2000; Pi & Lisman, 2008), so excessive CaMKII can weaken PP2A activity. Collectively, one explanation for impairment of NMDAR-dependent LTD is that the excessive αCaMKII can lower activity of PPs, thereby reduce GluA1-Ser845/Ser831 dephosphorylation and AMPAR internalization, and consequently impair LTD.

It has been shown that stargazin can be dephosphorylated by PP1 through Ca^2+^-PP2B-I1 pathway and form a ternary complex with APs to promote AMPAR internalization during NMDAR-dependent LTD (Matsuda et al., 2013; Tomita et al., 2003). Conversely, stargazin can be directly phosphorylated by activated CaMKII and bind to PSD-95 to immobilize AMPARs at synapses, which contributes to LTP (Bats et al., 2007; Opazo et al., 2010). In αCaMKII-F89G TG mice, more stargazin is expressed at the synaptic sites during NMDAR-dependent LTD. Therefore, another explanation for impairment of NMDAR-dependent LTD is that excessive CaMKII weakens AMPAR internalization through directly increasing stargazin phosphorylation and indirectly reducing stargazin dephosphorylation caused by lower PP1 activity, and finally impairs LTD.

## CONCLUSION

We have found that PTSD-susceptible mice exhibit the higher αCaMKII expression, and lower GluA1-Ser845/Ser831 dephosphorylation and AMPAR internalization in LA. Increasing αCaMKII leads to PTSD-like phenotypes such as impaired fear extinction and anxiety-like behaviors, and impairs LTD at T-LA synapses. Furthermore, diminished GluA1-Ser845/Ser831 dephosphorylation and AMPAR internalization were observed not only after impaired fear extinction *in vivo*, but also after attenuated NMDA-induced LTD in TG slices *in* vitro. Further data suggest that the impairment of NMDAR-dependent LTD is caused by the defective PPs activity and the excessive synaptic stargazin in αCaMKII-F89G TG mice. In summary, αCaMKII may be identified as a powerful regulator of the core symptoms of PTSD and LTD at T-LA synapses, and may be a key molecular determinant of PTSD.

## MATERIALS AND METHODS

### Animals

#### Biochemical Characterizations of αCaMKII-F89G TG mice

αCaMKII-F89G TG mice were donated by Dr. Tsien’s lab (Wang et al., 2003). Mutant αCaMKII-F89G was generated with silent mutation (i.e. replacing the Phe-89 with Gly in αCaMKII), so that the ATP-binding pocket of αCaMKII-F89G kinase was enlarged. To selectively block exogenous αCaMKII-F89G and leave endogenous αCaMKII intact, NM-PP1 was designed to fit only this enlarged pocket but not the unmodified pocket of native αCaMKII. By using αCaMKII promoter-driven construct, we were able to overexpress αCaMKII-F89G in the forebrain neurons. The αCaMKII-F89G could be rapidly and selectively manipulated in the mouse forebrain by intraperitoneal (i.p.) injection or noninvasive oral intake of 1-Naphthylmethyl (NM)-PP1. Specifically, a single i.p. injection of NM-PP1(16.57 ng/g) into freely behaving TG mice could completely suppress αCaMKII-F89G in the forebrain regions of TG mice within 15 minutes and the complete suppression could be maintained for 40 min. The oral intake (5 µM NM-PP1 in drinking water) could result in partial inhibition of αCaMKII-F89G in the TG mice by 6 h (no inhibition for the initial 3 h) and complete inhibition by 24 h. Bath application of 0.5 µM NM-PP1 in the slices of TG mice could inhibit αCaMKII-F89G but had no effect on native αCaMKII (Wang et al., 2003).

All experimental procedures were conducted according to Animals Act, 2006 (China) and approved by the Institutional Animal Care and Use Committee (IACUC approval ID #M09018) of the East China Normal University. All mice were male and 3-4 months old. C57BL/6J mice were used for Figure 1 and 3. αCaMKII-F89G transgenic mice and wild-type littermates were used for the rest of Figures. The mice were housed in 12 h light/12 h dark cycle (lights on at 7 a.m.) with free access to food and water.

### Behavior experiments

#### Behavioral profiling for identification of PTSD susceptible mice

We applied a behavioral profiling approach (Ardi et al., 2016) to identify PTSD susceptible mice in either underwater trauma (UWT)-exposed mice (G. Ritov, Boltyansky, & Richter-Levin, 2016) or 4 conditioned stimulus /unconditioned stimulus (4-CS/US)-exposed mice (Borghans & Homberg, 2015; Dębiec, Bush, & LeDoux, 2011; Fenster, Lebois, Ressler, & Suh, 2018; Ji et al., 2014a, 2014b; Mahan & Ressler, 2012; Mohammed R. Milad & Quirk, 2011; Radulovic et al., 2019).

In detail, the C57BL/6J mice were randomly divided into three groups: control group (n = 12), UWT-exposed group (n = 23) and 4-CS/US-exposed group (n = 23).

The control mice without any treatment were kept in home cages for 4 weeks. The UWT-exposed mice were individually allowed to swim freely for 5 s in a water-filled plastic tank, then submerged under water for 35 s using a metal net, next kept in their home cages for 4 weeks (Ardi et al., 2016; G. Ritov et al., 2016).

The 4-CS/US-exposed mice were individually placed in the chamber and allowed to explore the environment freely for 2 min, and then exposed to the conditioned stimulus (CS: 75 dB sound at 2800 Hz) for 30 s. At the last 2 s of tone stimulus, the unconditioned stimulus (US: 0.50 mA footshock, 2 s) was delivered. After 4-CS/US pairings with 2 min intertrial interval, mice were kept in the chamber for 2 min and then stayed in their home cages for 4 weeks.

Three groups were examined in the open field (OF) test, light/dark (LD) test, water zero maze (OM) test, fear conditioning and extinction experiments. The freezing behavior was monitored by Freeze Frame system (Coulbourn Instruments, USA).

We calculated six parameters: two parameters represent the level of locomotor activity and four parameters represent anxiety-like performances from the four experiments. To create the behavior profiles, firstly we referred to the performances of the control group as the behavior of the normal population and determined the distribution of values in the control group. Standard deviations were used to calculate the upper and lower “cut-off values” for each chosen parameter. Secondly, the performances of each mouse in the UWT-exposed group or 4-CS/US-exposed group were compared to the distribution curve of the control group. Each susceptible mouse must exhibit values that are under or above the lower and upper cut-off values in at least four out of the six parameters. “Cut-off values” of six parameters: the center time in the OF test, 560.32 ± 34.25 s; the time in the light box in the LD test, 788.60 ± 58.92 s; the time in the open arms in the OM test, 111.43 ± 8.88 s; the freezing percentage in the last day of cued fear extinction, 31.98% ± 3.91%; total distance in the OF test, 7059.99 ± 427.80 cm; total distance in the LD test, 9124.67 ± 220.50 cm.

#### Cued fear extinction

4 weeks after 4-CS/US parings or 24 h after the 1-CS/US paring, each mouse was placed into a novel chamber and monitored for 2 min (in the absence of the tone). For the recall test, the cued freezing responses to a 3 min tone (75 dB sound at 2800 Hz) without footshock were measured. Then, 4 cued fear extinctions trials were conducted like the recall test in the next 4 following days. Data were presented as the mean ± s.e.m. Two-way ANOVA was used for statistical analysis.

#### Open field

As described previously (Yan et al., 2015), briefly, each mouse was placed in an acrylic open-field chamber (27 cm long × 27 cm wide × 38 cm high) for 30 min. The amount of moving distance, the time in the center area, and the number of rearing were measured using a Tru-scan DigBahv-locomotion Activity Video Analysis System (Coulbourn Instruments, USA). Data were presented as the mean ± s.e.m. One-way ANOVA was used for statistical analysis in Fig. 1C, 2C and Student’s t-test in Fig. S2A.

#### Light/dark test

The box (27 cm long × 27 cm wide × 38 cm high) was divided into two equal zones -light zone and dark zone. The light zone was painted white and illuminated by the white light while the dark zone was painted black and not illuminated. These two zones were connected by a door in the middle divider. Mice could shuttle freely between two boxes. The total distance and the time stayed in light zone were delineated by the Tru-scan DigBahv-locomotion Activity Video Analysis System (Coulbourn Instruments, USA) for 30 min. Data were presented as the mean ± s.e.m. One-way ANOVA was used for statistical analysis in Fig. 1D and 2D.

#### Water-associated zero maze task

Experimental protocol and device were similar as described previously (Gilad Ritov & Richter-Levin, 2014).This device was composed of an annular platform and a plastic bucket. The annular platform was divided into four equal quadrants - two open arms and two closed arms. The plastic bucket was full of water for 40 cm deep. After 5 min habituation, mouse was put into one of the open arms facing the closed arm for 5 min. The time spent in the open arms and closed arms were measured by Any-maze system (USA, Stoelting). Data were presented as the mean ± s.e.m. One-way ANOVA was used for statistical analysis.

#### Elevated plus maze test

The apparatus consists of two opposed open arms (30 cm × 5 cm), two opposed closed arms (30 cm × 5 cm) and one open square (5 cm × 5 cm) in the center, which was elevated above the floor (50 cm). Each mouse was placed in the center of the plus maze with its face directing to an open arm and allowed to explore for 5 min. The time spent and moving distances in open and closed arms were automatically recorded by Any-maze system (USA, Stoelting). Data were presented as the mean ± s.e.m. One-way ANOVA was used for statistical analysis.

### Animal surgery

To elevate αCaMKII specifically in LA of C57BL/6J mice, we injected pAAV-TRE-αCaMKII-P2A-EGFP-CMV-rTA (AAV-αCaMKII) or pAAV-TRE-P2A-EGFP-CMV-rTA (AAV-control) virus (2.45 × 10^−12^ and 2.38 × 10^−12^ vector genomes/ml, respectively, Obio Technology, China) bilaterally into LA (AP, −1.60 mm; ML, ±3.35 mm; DV, −4.80 mm) of C57BL/6J mice. After the injection, mice were put back into home cages to recover for one month before experiments. AAV-αCaMKII mice were fed with doxycycline solution (1g/L in drinking water) to induce the virus expression throughout the behavior tests.

### Dendritic spine analysis

Dendritic spine analysis were performed as previously described (Ming et al., 2018). Briefly, mice were deeply anaesthetized and transcardially perfused. 200 μm coronal brain sections were cut and collected in 0.1 M PBS. LA neurons were loaded iontophoretically with a 5% Lucifer Yellow solution. Images of basal and apical dendrites of LA pyramidal neurons were scanned using a Leica SP2 confocal microscope at 63x under oil immersion. The number of spines per micrometer along the dendritic longitudinal axis was counted as spine density. Data were presented as the mean ± s.e.m. Student’s t-test was used for statistical analysis.

### Sensitivity to foot shock

This test was performed according to the methods as published (Duan, Zhou, Ma, Yin, & Cao, 2015). Mice were individually placed in the conditioning chamber to receive 1 s shocks of gradually increasing current intensity by an increment of 0.01 mA (flinching, 0.05-0.1 mA; vocalization, 0.1-0.2 mA; jumping, 0.45-0.6 mA) with 20 s intervals. The minimum current required to elicit flinching, vocalization and jumping in mice were measured. Data were presented as the mean ± s.e.m. Student’s t-test was used for statistical analysis.

### Amygdala slice electrophysiology

Protocols were similar as described previously (J. Kim et al., 2007; T. F. Ma et al., 2013). Mice (3-4 months old) were anaesthetized with sodium pentobarbital and sacrificed by decapitation. Whole brain coronal slices (370 μm thick for fEPSPs recording) containing the amygdala were cut using a vibroslicer (vibratome 3000) with the cold (4°C) and oxygenated (95% O_2_ /5% CO_2_) modified artificial cerebrospinal fluid (ACSF) containing (in mM): Choline choloride, 110; KCl, 2.5; CaCl_2_, 0.5; MgSO_4_, 7; NaHCO_3_, 25; NaH_2_PO_4_, 1.25; D-glucose, 25; pH 7.4. The slices were recovered in an incubation chamber with normal ACSF containing (in mM): NaCl, 119; CaCl_2_, 2.5; KCl, 2.5; MgSO_4_, 1.3; NaHCO_3_, 26.2; Na_2_HPO_4_, 1.0; D-glucose, 11, pH 7.4; 95% O_2_ and 5% CO_2_ for 60 min at 31°C, and then returned to room temperature for at least 1 h before recording.

### Field excitatory postsynaptic potential recording

A stimulating electrode was placed in the fibers from the internal capsule to activate the thalamic input to the lateral amygdala (T-LA) synapses. A recording electrode was positioned in LA to record field excitatory postsynaptic potential (fEPSP). Test responses were elicited at 0.033 Hz. After obtaining a stable baseline response for at least 15 min, LTP or LTD was induced. LTP was induced by applying 2 trains high frequency stimulation (100 Hz for 1 s) with 10 s interval or 3 trains theta burst stimulation (10 bursts delivered every 200 ms, each burst consisted of 4 pulses at 100 Hz) with 10 s interval. For LTD induction, the standard 1 Hz protocol (1 Hz for 15 min) and 3 Hz protocol (3 Hz for 5 min) were used. Depotentiation was induced by applying 2 trains of high frequency stimulation (100 Hz for 1s) with 10 s interval followed by the standard 1 Hz protocol (1 Hz for 15 min) after 20 min. For chemical-LTD induction, NMDA (Sigma, 30 μM in ACSF) was infused into the slice chamber for 3 min. Data were presented as the mean ± s.e.m. Student’s t-test (for comparing two different groups with Gaussian distribution) and one-way ANOVA followed by HSD post-hoc test with Bonferroni’s correction (for comparing more than two different groups) were used for statistical analysis.

### Proteins sample preparation

Combined with the previous protocol (Cui et al., 2011; Yin et al., 2013), synaptosomes were prepared as follows. LA tissues were homogenized in 1.5 ml homogenate-buffer (320 mM sucrose, 5 mM HEPES, pH 7.4) containing freshly added PMSF, PIC and PIC3. Homogenates were centrifuged at 500 g for 5 min to yield insoluble components. Then the supernatant fraction was collected and centrifuged at 10,000 g for 10 min to yield precipitation. The precipitation pellet was resuspended in 2 ml of 0.32 M sucrose, layered onto 2.25 ml of 0.8 M sucrose, and centrifuged at 98,000 g for 15 min using a swinging bucket rotor. Synaptosomes were collected from the 0.8 M sucrose layer and concentrated by centrifugation at 20,800 g for 45 min. Then the precipitation was resuspended in synaptosome lysis buffer (30 mM Tris (pH 8.5), 5 mM magnesium acetate, 8 M Urea, and 4% W/V CHAPS). For total proteins preparation, the LA areas were homogenized with RIPA buffer containing freshly added PMSF, PIC and PIC3 and lysed on ice for 30 min, centrifuged at 10,000 g at 4°C for 5 min, and total proteins were taken as supernatant. Then the protein samples were stored in a −80℃ freezer until used. Protein samples were quantified us by a Pierce BCA Protein Assay kit (Thermo Scientific) after which protein was stored at −20℃.

### Western blot

Each sample of protein (5 μg/lane) was separated by 10% SDS-PAGE (P40650, NCM Biotech) and separated at 120 V for 120 minutes. Then the separated proteins were transferred onto a polyvinylidene fluoride (PVDF) membrane. The PVDF membranes were blocked in blocking solution (5% skim milk and 1% BSA) at room temperature for 1h. A reversible Ponceau S staining of the membranes was done to normalize the relative amount of each protein on the membrane (just for synaptosomes). After washing with TBST buffer, the PVDF membranes were immunoblotted with following antibodies: GluA1 antibody (1:2,000, Santa Cruz), GluA2 antibody (1:2,000, Millipore), pGluA1-Ser845 antibody (1:500, Abcam), pGluA1-Ser831 antibody (1:500, Abcam), αCaMKII antibody (1:3,000, Abcam), p-αCaMKII-Thr286 antibody (1:20,000, Santa Cruz), βCaMKII antibody (1:2,000, Invitrogen), β-actin antibody (1:20,000, Sigma), GAPDH antibody (1:20,000, Proteintech), synapsin (SYP) antibody (1:2,000, Proteintech), TfR antibody (1:2,000, Abcam),Tubulin antibody (1:1,000, Millipore) at 4℃ for 12h. After washing with TBST buffer, the blots were reacted with an HRP-conjugated secondary antibody at room temperature for 1 hour. Band intensity on the blot was quantified by the ECL immunoblotting detection system (Bio-rad). Data were shown as mean ± s.e.m.. Statistical differences were analyzed using post hoc test with Bonferroni’s correction following one-way ANOVA.

### PP2A activity measurement

PP2A activity was measured by using immunoprecipitation phosphatase assay kit according to the manufacturer’s instructions (Catalog # 17-313, Millipore). Statistical differences were analyzed using post hoc test with Bonferroni’s correction following one-way ANOVA. Data were shown as mean ± s.e.m.

### PP2B activity measurement

The activity of calcineurin (PP2B) was assayed by using a calcineurin cellular activity assay kit (207007, Millipore) by following the manufacturer’s instructions. Statistical differences were analyzed using post hoc test with Bonferroni’s correction following one-way ANOVA. Data were shown as mean ± s.e.m.

### Statistical analysis

Statistical significance was assessed by one-way ANOVA, two-way ANOVA analysis of variance or two-tailed, unpaired and paired t-tests, where appropriate. Significant effects in analysis of variances were followed up with Bonferroni post-hoc tests. Results were considered significantly different when P < 0.05. All data were presented as means ± s.e.m. The detail information about statistical analysis was provided in legends.

## ACKNOWLEDGMENTS

We thank Dr. Xuechu Zhen and Dr. Liyong Li for their assistance with immunoblotting study, Dr. Kevan M. Shokat of UCSF for providing NM-PP1 inhibitor, and Dr. Bo Wang, Dr. Yihui Cui for their comments on the manuscript. This research was supported by fund from MOST China-Israel cooperation (No: 2016YFE0130500) and National Natural Science Fund of China (No: 31471077 and No: 31771177).

## AUTHOR CONTRIBUTIONS

S.A., J.W., X.Z., Y.D., J.L., D.W., H.Z., G.R.L., and X.C. designed the work; S.A., J.W., X.Z., J.L. and D.W. performed the acquisition of data for the work. S.A., J.W, X.Z., and X.C. analyzed and interpreted data; S.A., J.W, X.Z., G.R.L., and X.C. wrote the manuscript.

## ADDITIONAL INFORMATION

Supplementary information (Fig. S1, S2, S3A) accompanies this paper.

## Supplemental information

### Higher level of αCaMKII but normal morphology in LA of αCaMKII-F89G TG mice

First, we examined the αCaMKII expression level in LA of both αCaMKII-F89G TG and WT mice. Western blotting quantification revealed the expression of synaptic αCaMKII protein in LA of TG mice was 136% of WT littermates (Supplementary Fig. S1A, 1B, P < 0.05). Strikingly, the p-αCaMKII-Thr286 in LA of TG mice was 195% of WT littermates (P < 0.001). However, no obvious change in βCaMKII expression was observed in LA of TG mice (Supplementary Fig. S1A, B). Moreover, Nissl staining showed no detectable morphological abnormalities in LA of TG mice (Supplementary Fig. S1C). Normal shapes and architecture of dendritic spines could also be found in LA of TG mice (Supplementary Fig. S1D, S1E). These results suggest that the transgenic expression of αCaMKII-F89G increase αCaMKII expression in LA.

### Normal locomotor activity and acute pain threshold to footshock in αCaMKII-F89G TG Mice

Then, to investigate whether αCaMKII overexpression influences basal motor, exploratory behaviors and the foot shock sensitivity, we performed open field and pain threshold tests. No significant difference was observed between TG and WT mice in both locomotor activity (Fig. S2A, P > 0.05; Student’s t-test) and rearing behavior (Fig. S2A, P > 0.05), showing that TG mice exhibit normal locomotor activity and exploratory behavior. Moreover, we quantified the minimum current intensity of foot shock required to induce flinching, vocalizing and jumping in two groups of mice. There was also no significant difference in the threshold of current intensity to trigger flinching, vocalizing and jumping behaviors in TG mice and WT littermates (Fig. S2 B, P > 0.05). Taken all together, we can conclude that αCaMKII overexpression indeed impairs cued fear extinction.

**Fig S1.**
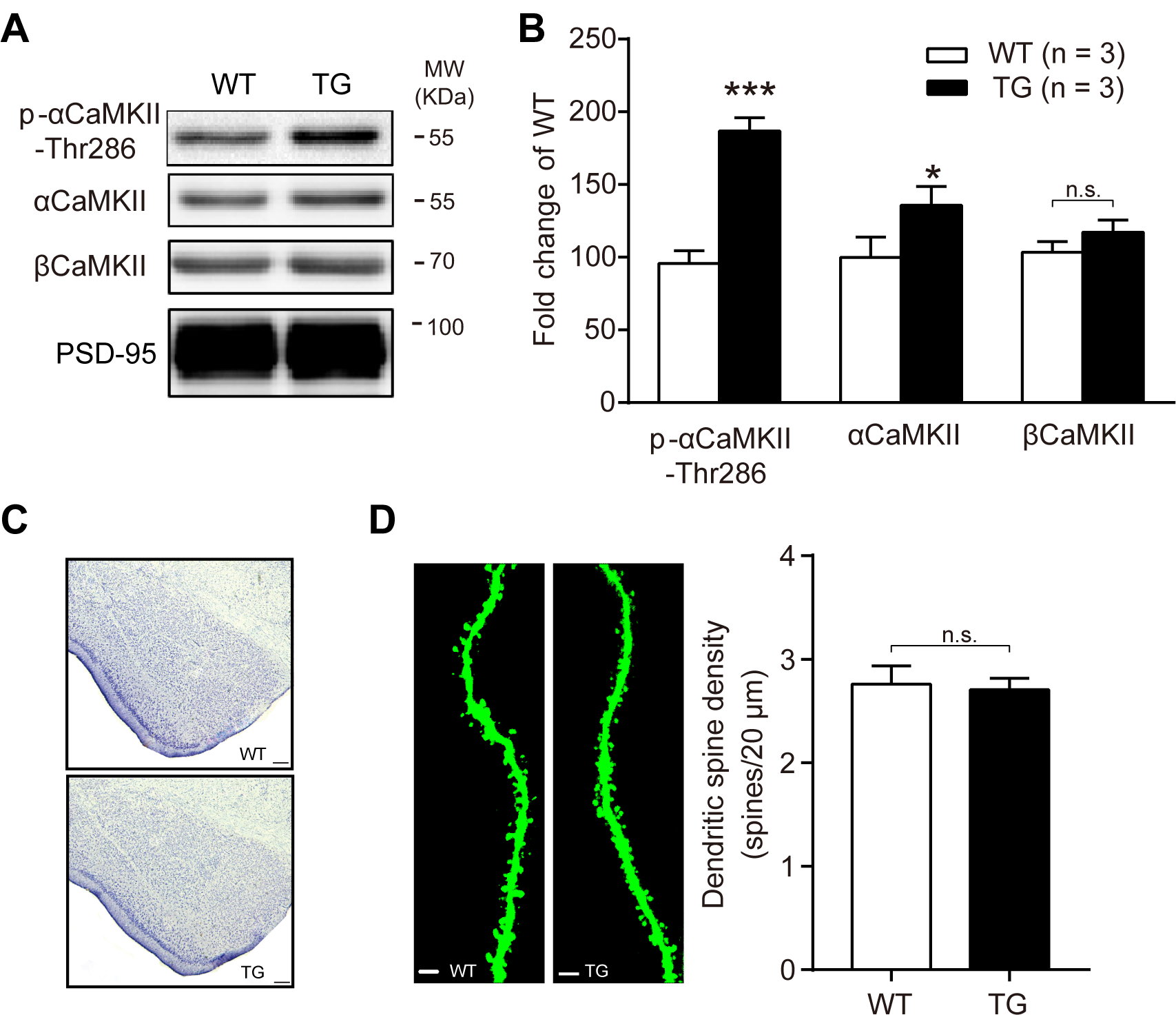
Higher level of αCaMKII but normal morphology in LA of αCaMKII-F89G TG mice. **(A)** Immunoblottings of αCaMKII protein in LA from WT and TG mice (p-αCaMKII-Thr286: p < 0.001; αCaMKII: p < 0.05; βCaMKII: p > 0.05). **(B)** Densitometric analysis shows a significantly higher expression of αCaMKII and p-αCaMKII-Thr286 in TG than that in WT mice. **(C)** Parts of Nissl stained coronal slices showing the amygdala of both WT and TG mice. Note no detectable morphological differences between WT and TG mice in the amygdala. Scale bars, 100 μm. **(D)** Dendritic spine of LA pyramidal neurons in WT and TG mice. Scale bar, 5 μm. The spine density (spines / 20 µm) was comparable between WT and TG mice (p > 0.05). Statistical differences were evaluated with Student’s t test, * P < 0.05, ** P < 0.01, *** P < 0.001. All data are shown as mean ± s.e.m.

**Fig S2.**
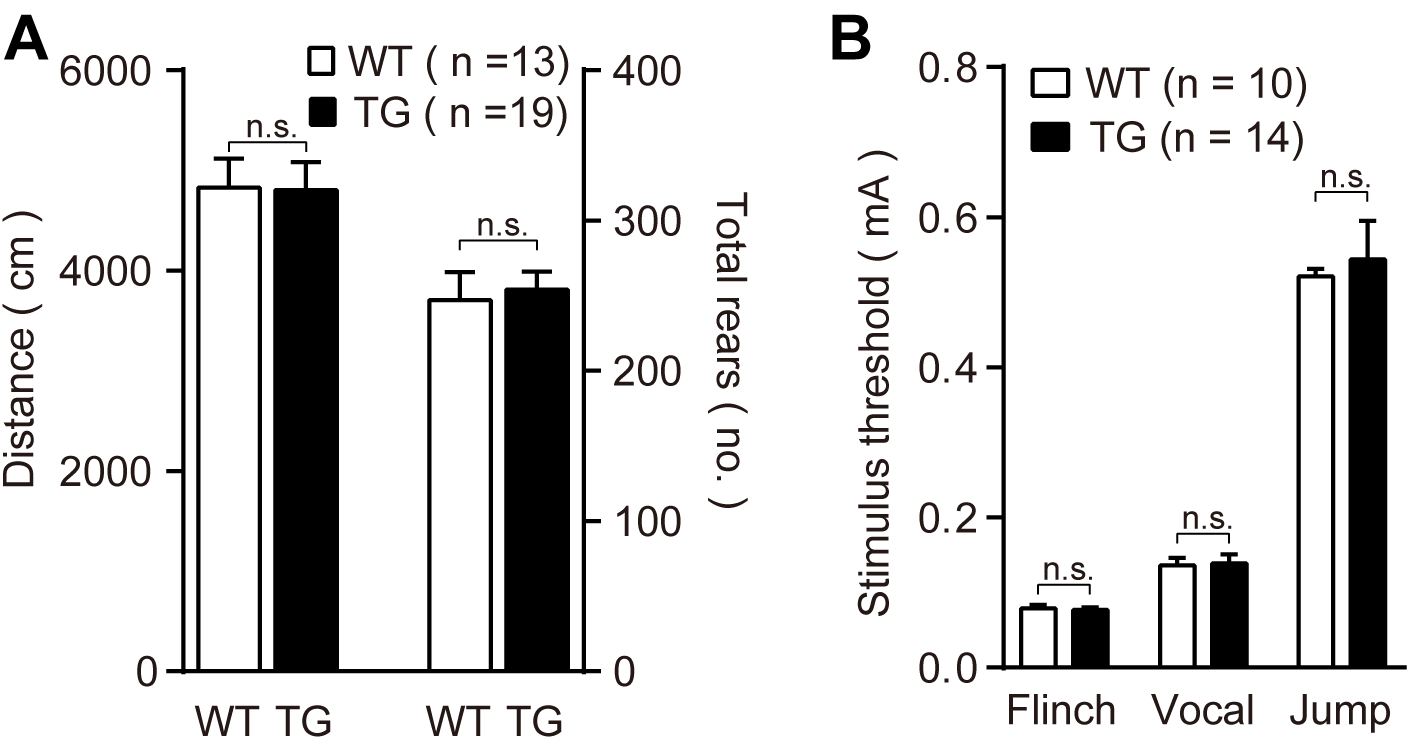
Normal locomotor activity and acute pain threshold to footshock in αCaMKII-F89G TG Mice. **(A)** Similar moving distance (P > 0.05) and rearing behavior (P > 0.05) in TG and WT mice during a 15 min of the open field test**. (B)** Normal pain sensitivity to an increasing electric footshock in TG mice (P > 0.05). All values are mean ± s.e.m. Statistical differences were evaluated with Student’s t-test.

**Fig S3.**
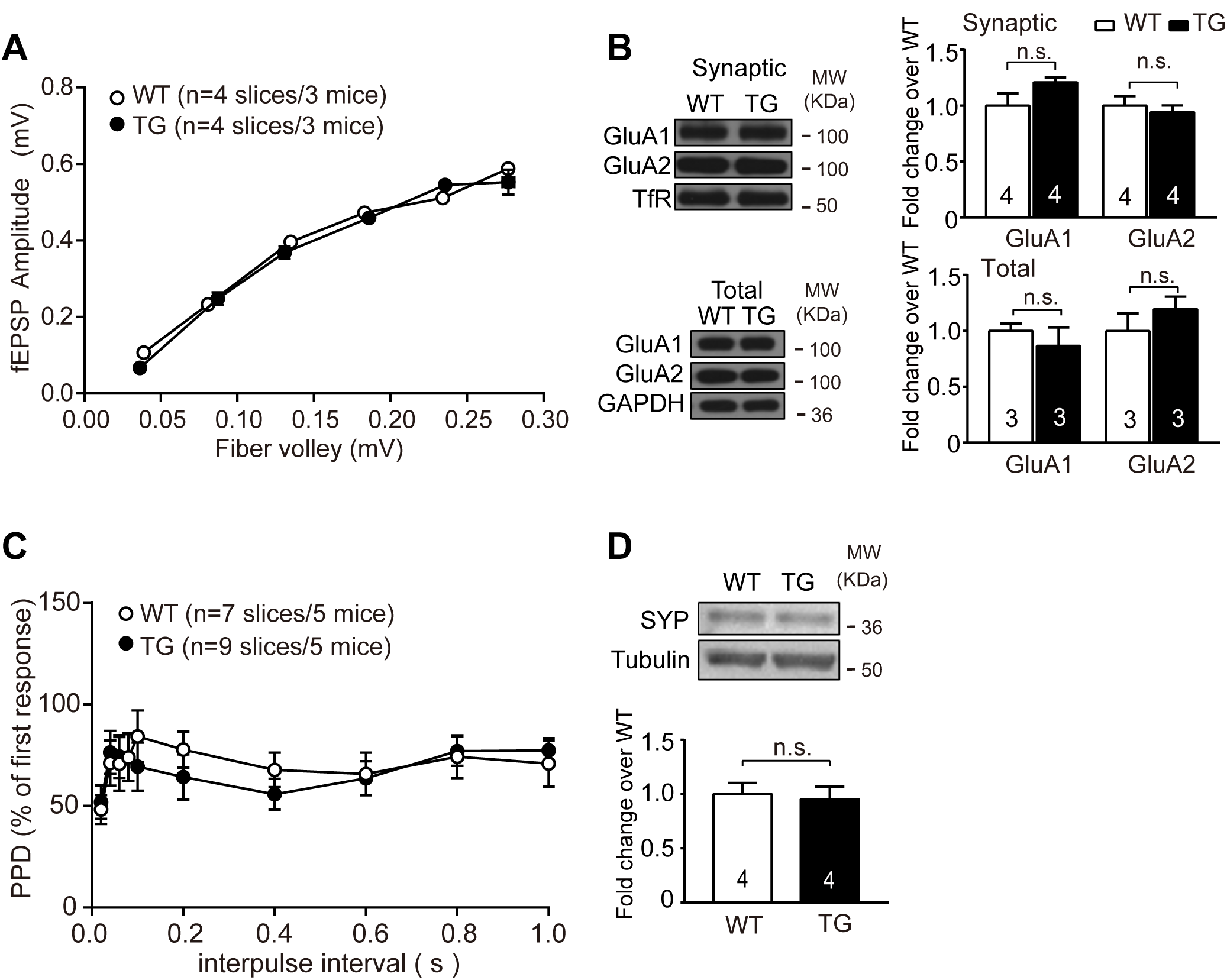
Normal basal synaptic transmission at T-LA synapses in αCaMKII-F89G TG mice. **(A)** No significant difference in the input/output curve at T-LA synapses between WT and TG slices (two-way ANOVA followed by multiple comparisons with Bonferroni’s correction). **(B)** Comparable synaptic or total GluA1/2 expression in LA of WT and TG slices (Statistical differences were evaluated with Student’s t-test). **(C)** Similar paired-pulse depression at different interpulse intervals in WT and TG amygdala slices (two-way ANOVA followed by multiple comparisons with Bonferroni’s correction). **(D)** Comparable expression levels of synapsin in LA of WT and TG amygdala slices (Statistical differences were evaluated with Student’s t-test). All data are shown as mean ± s.e.m.

